# An immobilization technique for long-term time-lapse imaging of explanted *Drosophila* tissues

**DOI:** 10.1101/2020.08.04.234864

**Authors:** Matthew P. Bostock, Anadika R. Prasad, Rita Chaouni, Alice C. Yuen, Rita Sousa-Nunes, Marc Amoyel, Vilaiwan M. Fernandes

## Abstract

Time-lapse imaging is an essential tool to study dynamic biological processes that cannot be discerned from fixed samples alone. However, imaging cell- and tissue-level processes in intact animals poses numerous challenges if the organism is opaque and/or motile. Explant cultures of intact tissues circumvent some of these challenges, but sample drift remains a considerable obstacle. We employed a simple yet effective technique to immobilize tissues in medium-bathed agarose. We applied this technique to study multiple *Drosophila* tissues from first-instar larvae to adult stages in various orientations and with no evidence of anisotropic pressure or stress damage. Using this method, we were able to image fine features for up to 18 hours and make novel observations. Specifically, we report that fibers characteristic of quiescent neuroblasts are inherited by their basal daughters during reactivation; that the lamina in the developing visual system is assembled roughly 2-3 columns at a time; that lamina glia positions are dynamic during development; and that the nuclear envelopes of adult testis cyst stem cells do not break down completely during mitosis. In all, we demonstrate that our protocol is well-suited for tissue immobilization and long-term live imaging, enabling new insights into tissue and cell dynamics in *Drosophila*.

## Introduction

Live imaging is a powerful tool to elucidate mechanistic and temporal aspects of intricate biological processes. Dynamic processes such as cell migration, protein localization, axon pathfinding and branching morphogenesis are described poorly in fixed tissue, whereas live imaging can reveal features within these processes with exquisite temporal resolution (Besson *et al*., 2015; Rabinovich *et al*., 2015; Chen *et al*., 2016). This approach has seen dramatic improvements with 2-photon and light sheet microscopy due to the increased depth of access and diminished phototoxicity (Huisken and Stainier, 2009; Nickerson *et al*., 2013; Ichikawa *et al*., 2014). Furthermore, developments in sample preparation for *in vivo* and *ex vivo* imaging as well as in advanced computational analyses have increased accessibility to investigations of dynamic processes (Ritsma *et al.*, 2014; Speder and Brand, 2014; Rabinovich *et al.*, 2015; Martin *et al.*, 2018). Notwithstanding, a major obstacle with live imaging is sample drift, which results in a structure of interest moving out of focus. This can pose challenges to image analysis of dynamic processes.

Sample drift has been combatted by using coverslips or glass slides coated with adhesive extracellular matrix proteins such as fibronectin or collagen to physically immobilize the tissue of interest. However, these steps may exert extraneous anisotropic physical stress on the sample and affect developmental mechanisms, cause injuries to fragile tissues and therefore significantly reduce imaging time (Savoian and Rieder, 2002; Siller *et al*., 2005; Lerit *et al*., 2014; Rabinovich *et al*., 2015). Solutions to these problems have included placing explants in agarose wells (Rabinovich *et al*., 2015) but without being held in place, they still move. Although there are computational algorithms that can account for sample drift, they are often slow and can result in discontinuities between frames thus decreasing confidence in the image (Parslow *et al*., 2014).

Live imaging has been applied to many systems but here we focus on *Drosophila melanogaster*, whose genetic tractability makes it an outstanding model to image dynamic cellular processes. The *Drosophila* embryo was one of the earliest animal systems imaged live, due to being translucent and immobile up to late stages. Dechorionated live embryos can be imaged by gluing to a coverslip and covering with halocarbon oil to minimize dehydration (Cavey and Lecuit, 2008; Parton *et al*., 2010). Live imaging of the *Drosophila* embryo has been used widely to elucidate nuclear and cytoplasmic behaviors in the preblastodermal embryo (Foe and Alberts, 1983; Baker *et al*., 1993), epithelial adhesion during dorsal closure (Jacinto *et al*., 2000; Kiehart *et al*., 2000), germ cell migration (Sano *et al*., 2005), neuroblast divisions (Kaltschmidt *et al*., 2000) and mechanisms of salivary gland formation (Sanchez-Corrales *et al*., 2018) among many others. Beyond the embryo, *in vivo* live imaging becomes challenging since larvae and adults move continuously and have opaque cuticles which scatter light (Aldaz *et al*., 2010; Rabinovich *et al*., 2015; Bell, 2017). Calcium oscillations across the blood-brain barrier have been imaged through the thinner cuticle of very young larvae, reasonably steadied between coverslip and culture dish (Spéder and Brand, 2014). Notwithstanding, while this methodology was apt for capturing relatively large-scale inter-cellular calcium wave propagation, the considerable drift remaining is not suited to visualize finer (sub)cellular events. Similarly, although larvae and pupae have been imaged live, the need to strike a balance between phototoxicity and image-acquisition rates often mean that some dynamic processes are hard to capture (Bosveld *et al*., 2012; Ghannad-Rezaie *et al*., 2012; Heemskerk *et al*., 2014; Tsao *et al*., 2016; Dye *et al*., 2017). In adults, live imaging can be performed through windows cut out of the cuticles of immobilized animals (Fiala *et al*., 2002; Seelig *et al*., 2010; Martin *et al*., 2018; Aimon *et al*., 2019) but feasibility of this approach depends on the accessibility of the tissue of interest.

An alternative to *in vivo* imaging is to image tissues in culture. Initially, explanted tissues were imaged to study processes over short periods of time (*i.e.* minutes to hours) such as cell cycle progression and oriented cell divisions, epithelial cell packing, intracellular protein movements and secretion (Siller *et al*., 2005; Farhadifar *et al*., 2007; Siller and Doe, 2008; Aldaz *et al*., 2010; Mao *et al*., 2011; Lerit *et al*., 2014). More recently, live imaging of cultured explants has been extended to processes that unfold over several hours such as morphogenesis of pigment cells during pupal eye development (Hellerman *et al*., 2015), cell migration (Prasad *et al*., 2015; Chen *et al*., 2016; Barlan *et al*., 2017), neuronal remodeling (Rabinovich *et al*., 2015), growth cone dynamics (Özel *et al*., 2015; Akin and Zipursky, 2016), and spermatogonial stem cell dynamics in their niche (Sheng and Matunis, 2011).

Different culture media compositions have been applied to long-term live imaging of explanted *Drosophila* tissues. The most commonly used is Schneider’s Insect medium (Echalier, 1997). Echalier’s D-22 medium (Siller *et al.*, 2005; Lee *et al*., 2006), Shield’s and Sang’s M3 medium (Aldaz *et al*., 2010) and Grace’s Insect Culture medium have also been employed (Dye *et al*., 2017). These media are often supplemented with exogenous growth supporting components such as insulin, fetal bovine serum, fly extract, larval fat bodies, ascorbic acid and/or 20-hydroxy-ecdysone (20E) to optimize culture conditions. Supplement requirements vary with the tissues being imaged (Parton *et al*., 2010) and there are conflicting opinions regarding supplements for the same tissue. For example, some studies report that the addition of fly extract is essential to support imaginal disc growth *ex vivo* (Wyss, 1982; Zartman *et al*., 2013; Restrepo *et al*., 2016) whereas others demonstrated that fly extract had no effect on disc growth and in fact caused aberrant calcium oscillations in cultured wing discs (Tsao *et al*., 2016; Balaji *et al*., 2017). Similarly, larval fat bodies were found to be vital to maintain neuroblast divisions *ex vivo* (Siller *et al*., 2005; Cabernard and Doe, 2013) but others either found them dispensable for neural proliferation in the young larval central nervous system (CNS) or inhibitory of early pupal CNS development (Sousa-Nunes *et al*., 2011; Rabinovich *et al*., 2015). Lastly, addition of 20E and insulin to culture medium aimed at supporting imaginal disc growth has also been debated. Absence of insulin and presence of 20E has been reported to enable disc growth *ex vivo* (Aldaz *et al*., 2010; Dye *et al*., 2017) although other studies suggest that insulin is necessary (Restrepo *et al*., 2016; Tsao *et al*., 2016) but that 20E impairs disc development (Tsao *et al*., 2016). Differences in tissue responses to these supplements might be attributed to the specific stage and/or basal medium being used. For example, Zartman *et al*. demonstrated that cells derived from wing discs proliferated to a greater extent when insulin was added to Schneider’s Insect Medium but not M3 medium (Zartman *et al*., 2013).

Here, we present a simple protocol for culturing and imaging *Drosophila* larval and adult tissues *ex vivo*. We use Schneider’s Insect Medium along with relatively few growth supplements and immobilize samples in low gelling temperature agarose, an adaptation of the method commonly used to immobilize zebrafish embryos or larvae for live imaging (Distel and Koster, 2007). In this way, the explanted tissue is held in place without imposing anisotropic physical stress on it. Moreover, this technique allows tissues to be held in any orientation, independent of shape and center of gravity, rendering imaging of fine features readily accessible. We have used this approach successfully to image the migration of glial cells and neurons in the *Drosophila* brain during the third larval instar (Chen *et al*., 2016; Rossi and Fernandes, 2018). Here, we validate our protocol in multiple tissues from multiple developmental stages and report new biological observations for the first time. Specifically, we followed neuroblast divisions not only in the commonly imaged wandering third larval instar (wL3) brain but also as they reactivate from quiescence during the first and second larval instars (L1 and L2); we captured glial and neuronal migration in the optic lobe, assembly of lamina columns and eye-antennal disc eversion; and we imaged cyst stem cell mitoses in adult testes. Overall, this is an inexpensive and simple method to carry out live imaging experiments to broaden understanding of cell and tissue dynamics in *Drosophila*.

## Methods and Materials

### Fly husbandry and stocks

Fly strains and crosses were raised on standard cornmeal food at 25 °C, except for the sparse labelling of epithelial and marginal glia (*dpp*>*FlexAmp*), which was raised at 29 °C.

The Following genotypes were used in this study: *{yw; gcm-GAL4/CyO;}* Bloomington Drosophila Stock Center (BDSC) #3554, *{;; UAS-CD8::GFP/TM6B}* BDSC #5130, *{;; UAS-nls::GFP/TM6B}* BDSC #4776, *{; 13xLexAop-6xmcherry/CyO;}* BDSC #52271, *{yw, UAS-FLP; GAL80*^*ts*^*/CyO; Act*>*y+*>*lexA, lexAop-myr::GFP /TM6B} (FlexAmp)* (Bertet *et al*., 2014), *{;; dpp-GAL4/TM6B}* BDSC #7007, *{; E-Cad-E-Cad::GFP;}* (Huang *et al*., 2009), *{;; R27G05-LexA/TM6C}* (Tan *et al*., 2015), *{; ubi-GFP::CAAX;}* Drosophila Genomics Resource Center (DGRC) *#109830, {; His2av::EGFP/SM6a;}* BDSC #24163, *{; Tj-GAL4;}* DGRC #104055, *{; grh-GAL4;}* (Chell and Brand, 2010), *{;; UAS-syn21-GFP-p10}* (Pfeiffer *et al*., 2012), *{;; UAS-CD4-tdTomato}* a gift from D, Williams.

### Explant culture medium

Explant culture medium consisted of Schneider’s Insect medium (Sigma #S0146) supplemented with 1.25 mg/ml human insulin (Sigma #I9278), 1 % Penicillin-Streptomycin (Sigma #P4333) and 10 % Fetal Bovine Serum (Sigma #F2442) stored at 4 °C and used within a month of preparation. See Supplementary Materials for detailed protocol with suggested volumes.

### Preparation of low gelling temperature agarose

2 % low-gelling temperature agarose (Sigma #A9414) was prepared in phosphate buffered saline. These were cut into ∼ 0.5 cm^3^ pieces and stored in distilled water at 4 °C. The agarose was deionized by changing the water each day for 5 days before use. See Supplementary Materials for a detailed protocol with suggested volumes.

### Dissections

Forceps, dissection pads, pipettes, falcon tubes and working areas were wiped down with 70 % ethanol before use. Dissection of L1, L3 and adult tissues was carried out in cold culture medium. Dissections of L1 CNSs were performed using forceps to hold down the posterior end of the larva and a tungsten needle to slowly rip open the larval cuticle, and then lightly pull on the mouth hooks to extract the CNS. CNSs were left attached to mouth hooks via the esophagus, as well as to surrounding fat tissue and imaginal discs to avoid damage. Dissections of L3 CNSs were performed with a pair of forceps, used to rip and remove the larval cuticle, and sever the CNS from the midgut. Fat tissue and imaginal discs were removed, leaving only mouth hooks attached to the CNS via the esophagus. For L1-L3 CNS imaging esophageal muscles were crushed to cease unwanted contractions. For adult testes, flies were dissected 0-3 days post-eclosion with careful removal of the ejaculatory duct and accessory glands of the male gonad leaving each testis intact with its connecting seminal vesicle.

### Tissue immobilization in agarose

Deionized agarose (see above) was melted in a microwave for approximately 20-30 seconds (per 0.5cm^3^ cube of 2 % low-gelling temperature agarose) and diluted to 0.4 % in culture medium heated to 42 °C using a programmable heating block. The temperature was then lowered to 34 °C before being added to coat the bottom of untreated 35 x 10 mm petri dishes (Thermo #171099). A single explant was placed in each dish and maneuvered to the center to be oriented using forceps. To maneuver the tissue, forceps were used to move the viscous agarose rather than the tissue itself. Once the desired orientation was achieved, the forceps were gently withdrawn from the agarose, which held the tissue in place due to its viscosity. All movements and orientations of the tissue were achieved within 5 mins of placing the brain in the agarose so as not to disrupt its setting. The agarose was left to solidify for 10 mins after which cold culture medium was added. See Supplementary Materials for a detailed protocol with suggested volumes.

### Imaging and image processing

We used an upright microscope set-up (Olympus FV1000MPE multi-photon laser scanning microscope or Zeiss 880, both with Spectra-Physics MaiTai DeepSee 2-photon lasers) with water-immersion lens (Olympus XLPLN 25X WMP2 or Zeiss Plan-Apochromat 20X). The fluorophores used were all GFP or RFP derivatives, therefore the excitation wavelength was tuned between 925nm and 935 nm. Laser power never exceeded 15 %. Zen Blue (Zeiss) and ImageJ software was used to analyze movies. The Bleach Correction and Manual Tracking plugins were used to correct photobleaching and to track cells. The Correct 3D Drift plugin was used to correct for movements caused by tissue contraction. Adobe Photoshop (v21.1.3) and Adobe Premiere Pro (v14.2) were used to annotate and edit movies. Figures were generated using Adobe Illustrator (v24.1.3).

## Results

### Neuroblast divisions in the L3 central brain

The late larval CNS has been used extensively to study the biology of neural stem cells, called neuroblasts in *Drosophila*. Neuroblasts generate a vast number of diverse neuronal and glial cell subtypes which are critical for neural function. So-called type I neuroblasts are the most abundant and are found throughout the CNS (Figure 1). They divide asymmetrically to self-renew and generate a transit-amplifying progenitor called ganglion mother cell (GMC) (reviewed by Sousa-Nunes and Hirth, 2016). The GMC then undergoes a terminal division to produce two neuronal and/or glial progeny whereas the self-renewed neuroblast continues to proliferate. To compare our protocol to existing strategies for visualizing neuroblast dynamics, we imaged divisions in the central brain from animals dissected at the wL3 stage (Figure 1). To visualize chromatin we used *His2Av::eGFP*, a histone variant fused to an enhanced green fluorescent protein (eGFP); larval neuroblasts were identified as large (∼10-15 μm diameter) superficial cells (Truman and Bate, 1988). As expected, these cells underwent a self-renewing division to generate a neuroblast and a GMC (Movie 1). Between divisions, neuroblasts grew in size and their cell-cycle time was ∼90 minutes, consistent with other reports (Cabernard and Doe, 2013; Homem *et al*., 2013). We also observed GMC divisions (Movie 1). Although it has been reported that larval fat bodies are essential for sustaining neuroblast divisions *ex vivo* (Siller *et al*., 2005; Cabernard and Doe, 2013), we found them to be dispensable. In summary, neural progenitor divisions proceeded as expected demonstrating that our culture medium and immobilization technique can support them.

**Figure 1.**
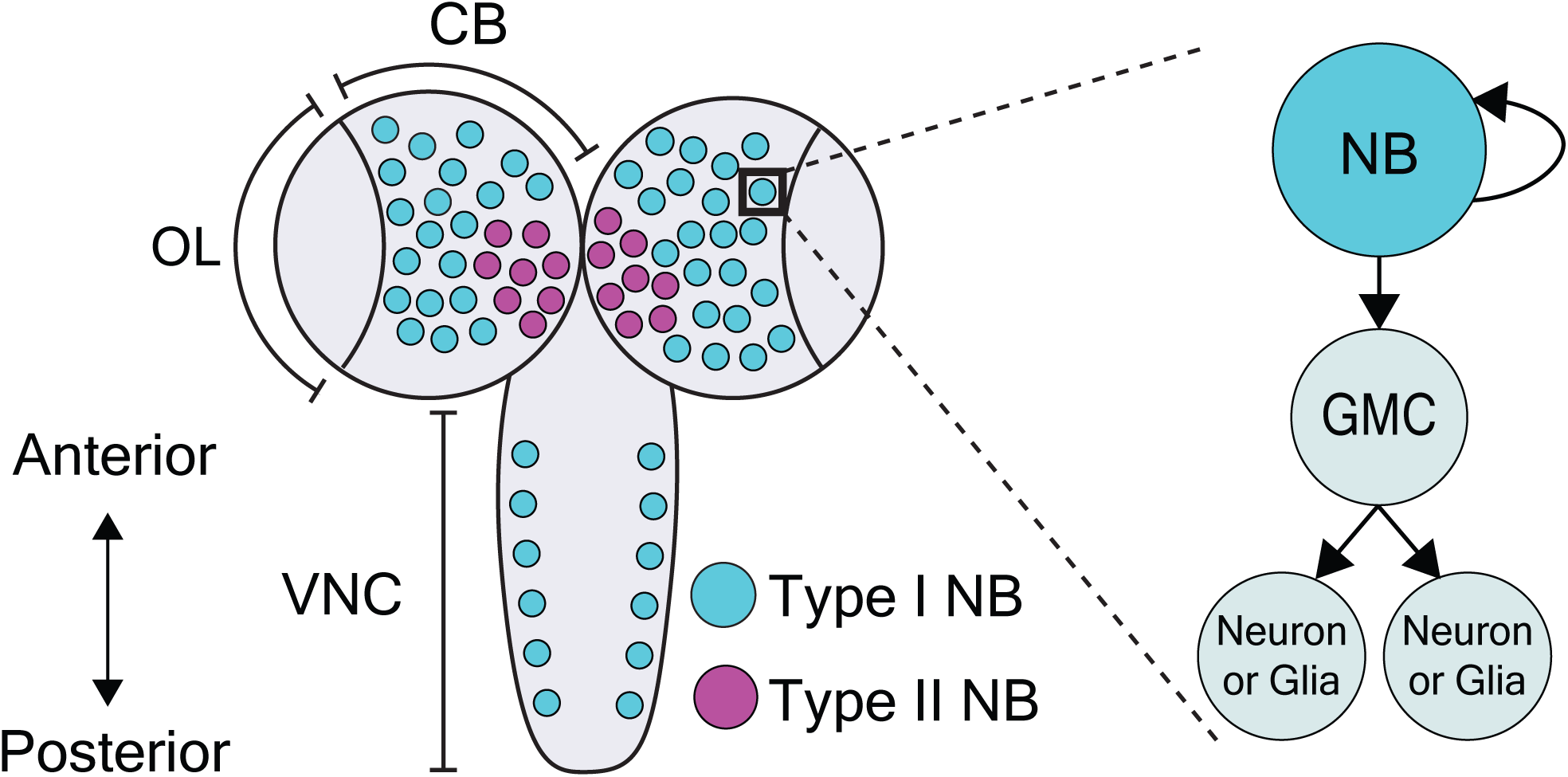
Dorsal view schematic of the third-instar larval central nervous system. The L3 CNS is made up of the optic lobe (OL), the central brain (CB) and the ventral nerve cord (VNC) (Sousa-Nunes and Hirth, 2016). Type I neuroblasts (blue), which divide to self-renew and generate a GMC, are most abundant and are present throughout the CNS. GMCs divide symmetrically in size to generate two differentiating neuronal or glial progeny. Type II neuroblasts (magenta), defined by generation of two types of transit-amplifying progenitors (intermediate neural precursors and GMCs) consist of eight paired lineages found in the dorsoposterior regions of the CB. The black box indicates the region selected for live imaging, which contains only type I neuroblasts.

### Neuroblast reactivation

In contrast to many studies employing live imaging of neuroblasts from L3 CNSs, none as yet report imaging of neuroblast divisions in the more fragile first or second larval instars (L1/L2). Nonetheless, these earlier stages include specific processes of interest. By the end of embryogenesis, most neuroblasts enter a state of reversible cell cycle arrest termed quiescence (Truman and Bate, 1988; Valcourt *et al*., 2012). Following larval hatching and feeding, neuroblasts exit quiescence (reactivate) in an anteroposterior order (Truman and Bate, 1988; Britton and Edgar, 1998; Chell and Brand, 2010; Sousa-Nunes *et al*., 2011). Neuroblasts are relatively large when actively proliferating (10-15 µm diameter) (Chell and Brand, 2010; Sousa-Nunes *et al*., 2011) and are devoid of morphological polarity despite extensive molecular asymmetries during mitosis. In contrast, quiescent neuroblasts are much smaller (∼4 µm diameter) and morphologically polarized, projecting a basal fiber into the neuropil (Figure 2). This morphology is reminiscent of that of vertebrate radial glia (Weissman *et al*., 2003) and renders quiescent neuroblasts morphologically indistinguishable from adjacent neurons (Figure 2). As they reactivate, neuroblasts enlarge and lose their fiber. We wondered whether fibers would be severed or retracted during neuroblast reactivation and endeavored to image this process live.

**Figure 2.**
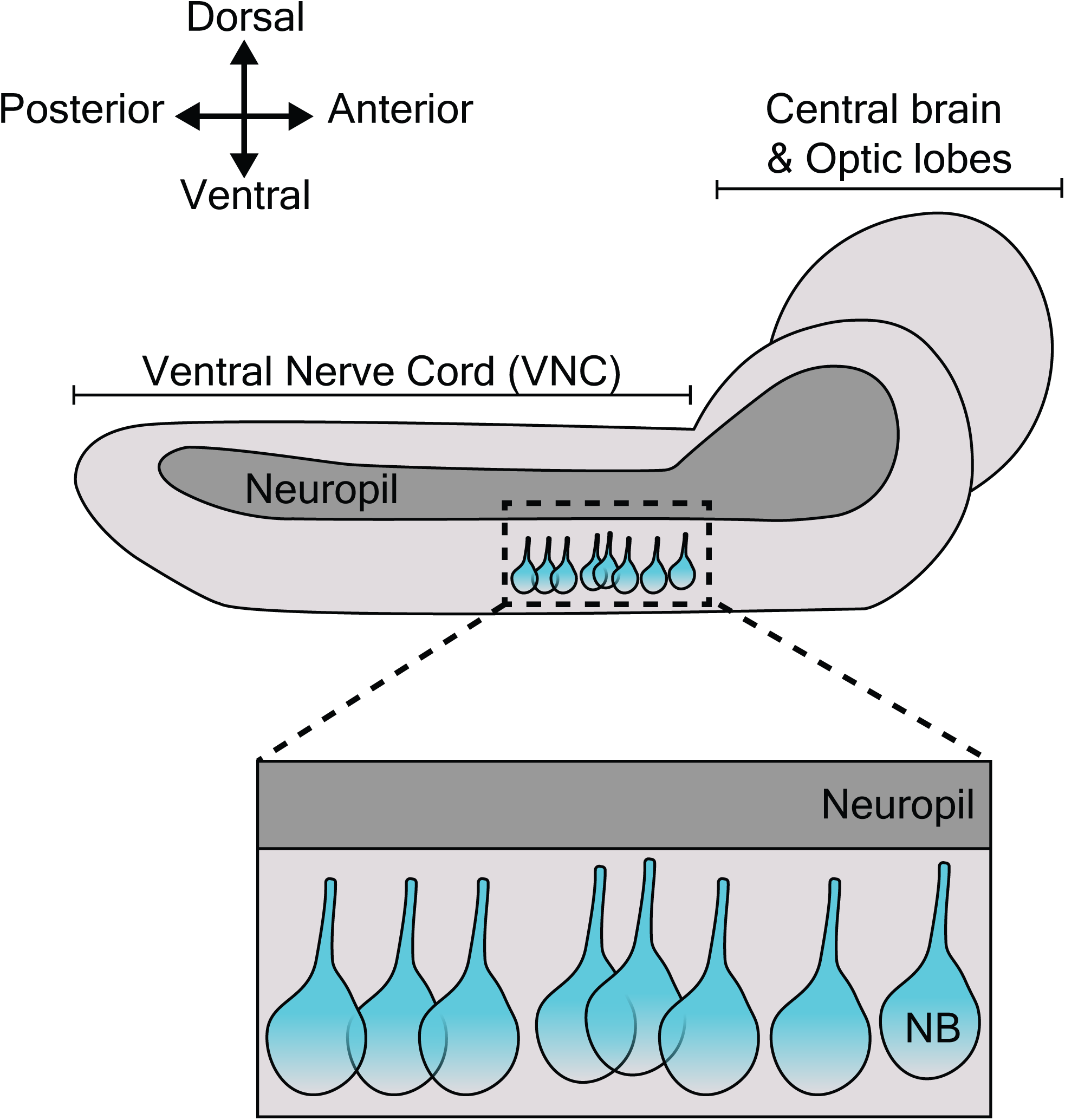
Lateral view schematic of L1/L2 CNS depicting quiescent neuroblast morphology. Neuroblasts, depicted in blue, are shown within the thoracic region of the ventral nerve chord (VNC, dashed box), the region selected for live imaging. Neuroblast (NB) cell bodies are situated on the ventral side of the VNC. During quiescence, they extend a fiber towards the neuropil. Upon reactivation, which occurs roughly 24 h after larval hatching for neuroblasts in the thoracic VNC, neuroblasts lose their fiber, which marks a relatively early morphogenetic event we sought to capture through our live imaging protocol.

Young larval CNSs are more susceptible to mechanical stress than later ones, including to forces exerted by laminin or poly-L-lysine-coated surfaces (our own observations). Immobilizing L1 or L2 brains in this way invariably resulted in CNS rupture. While L1 and L2 CNSs did not rupture when immobilized under a fibrin clot (Lerit *et al*., 2014), we were unable to orient them at will to visualize neuroblasts clearly using this method (data not shown). The agarose-based immobilization approach described here proved sufficiently gentle and allowed for the desired orientation, enabling documentation of neuroblast reactivation for the first time. *grainyhead (grh)-GAL4*, expressed in a subset of neuroblasts (Chell and Brand, 2010) was used to drive expression of *UAS-Syn21-GFP-p10*, a translationally enhanced GFP reporter (Pfeiffer *et al*., 2012). Several ventral nerve cord neuroblasts were observed reactivating and undergoing mitosis over the course of 17 hours (Movie 2). Mitoses were clearly recognizable by cells rounding prior to dividing into one larger apical daughter (renewed neuroblast) and one smaller basal daughter (the GMC). Neuroblast and GMC divisions continued after reactivation in a few cases (n = 6) indicating favorable conditions.

To our surprise, we found that neuroblasts retained their fiber throughout the first post-reactivation division and that the fiber was inherited by the first post-reactivation GMC (Movie 3, n = 19) and then by GMC progeny (n = 5). We could not discern whether GMC fibers were asymmetrically inherited by one of the ganglion cell daughters (usually neurons) or split and inherited by both. Asymmetric basal fiber inheritance has been described for zebrafish, rodent and human embryonic/fetal neural progenitors. Intriguingly, in contrast to what we observed in *Drosophila*, in those models it was generally the self-renewing progenitor that inherited the fiber (Weissman *et al*., 2003; Konno *et al*., 2008; Alexandre *et al*., 2010; Hansen *et al*., 2010; Shitamukai *et al*., 2011) although, on occasion, asymmetric inheritance by neuronal progeny was observed (Miyata *et al*., 2001; Konno *et al*., 2008), as was fiber splitting and seemingly symmetric inheritance by both daughter cells (Konno *et al*., 2008). We speculate that fiber inheritance by GMCs and then neuronal progeny could be a mechanism to quickly develop neurites, especially important for a fast-developing organism like *Drosophila.*

The above demonstrates that our protocol is well-suited to immobilize early larval brains even in the generally unstable side orientation for long-term neuroblast imaging, including observation of the first post-reactivation division and GMC fiber inheritance from quiescent neuroblasts, which has not been reported before.

### Lamina development in the L3 optic lobes

Next, we turned our attention to the developing L3 optic lobe, specifically focusing on the developing lamina. The lamina arises from a crescent-shaped neuroepithelium called the outer proliferation center (OPC), which is located at the surface of the optic lobe. The lateral edge of the OPC folds to form a structure called the lamina furrow (LF), from which lamina precursor cells (LPCs) are generated (Figure 3A and B). Photoreceptors from the eye disc grow their axons through the optic stalk and into the optic lobe where they defasciculate and contact the LF along the dorsoventral length of the OPC crescent. R1-R6 photoreceptors terminate their growth cones at the level of the LF (Figure 3B). Photoreceptors deliver Hedgehog through their axons and induce LPC formation from LF neuroepithelial cells (Figure 3A) (Huang and Kunes, 1996; Huang *et al*., 1998b, 1998a). LPCs then associate with photoreceptor axons to form columns before differentiating into lamina neurons (Huang and Kunes, 1996; Huang *et al.*, 1998b, 1998a; Umetsu *et al.*, 2006; Fernandes *et al.*, 2017).

**Figure 3.**
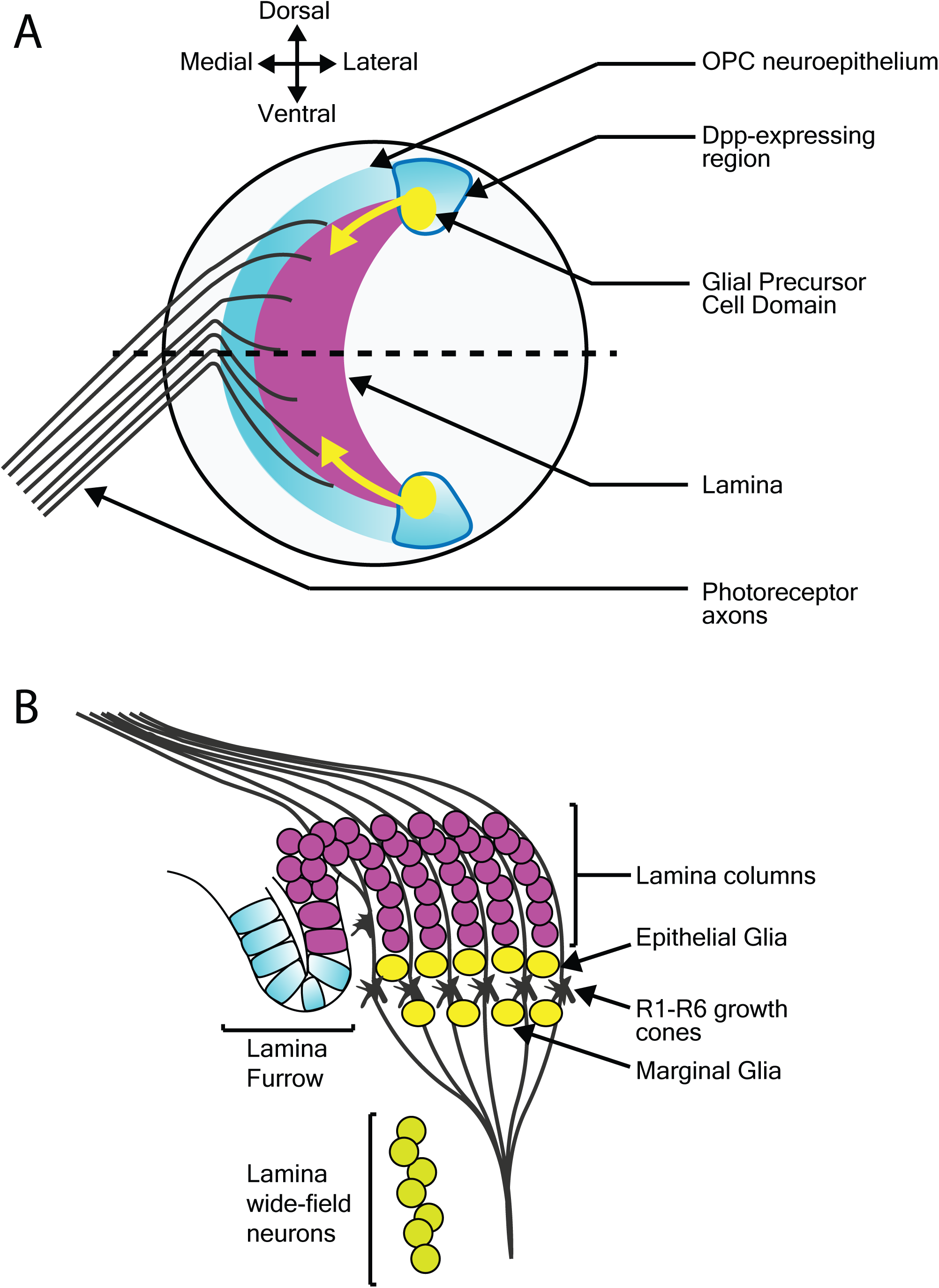
Migration of epithelial and marginal glia, and lamina wide-field neurons (Lawf). **A:** Schematic of the lateral view of the third larval instar optic lobe. Developing photoreceptors project to a fold in the outer proliferation center neuroepithelium called the lamina furrow and induce lamina formation. **B:** Diagram of a cross-section along/at the level of the dotted line in A. The developing lamina forms in characteristic columns. Seven lamina precursor cells are incorporated into each column. Epithelial and marginal glia migrate above and below photoreceptor growth cones. Lawf neurons share the same progenitors as epithelial and marginal glia; they migrate from their point of origin at the tips of the lamina to the medulla, where they stop immediately adjacent to the neuropil.

To visualize lamina development we used *E-Cadherin::GFP* (*E-Cad::GFP*), which localizes to epithelial adherens junctions, together with the lamina-specific *(R27G05-GAL4)* expression of cytoplasmic *mCherry* (Movie 4). Photoreceptor axons and the lamina furrow showed enriched *E-Cad::GFP* expression (Movie 4). The lamina grew dramatically over the course of ∼18 hours (Movie 4). Interestingly, throughout this growth the lamina furrow remained relatively stable (Movie 4 – asterisk), with lamina growth displacing older lamina columns posteriorly (Movie 4). This is in contrast to previous assumptions based on fixed images that the lamina furrow moved similarly to the morphogenetic furrow in the eye imaginal disc (Selleck and Steller, 1991; Huang and Kunes, 1998a) and implies a different process by which LPCs are generated from the neuroepithelium.

The use of cytoplasmic mCherry prevented us from distinguishing individual LPCs and their incorporation into columns. We therefore switched to nuclear GFP (*UAS-nlsGFP*) driven by *glial cells missing (gcm)-GAL4*, which marks LPCs and lamina glia (Figure 3 and Movie 5). We manually tracked LPCs as they exited the LF through to incorporation into columns. Rather than column assembly progressing one column at a time, we observed that LPCs incorporated into the first 2-3 columns simultaneously, suggesting that multiple young columns are assembled together (Movie 5). This was surprising since it was generally assumed that the lamina is built one row of columns at a time (Umetsu *et al.*, 2006; Sugie *et al*., 2010; Sato *et al*., 2013).

### Glial and neuronal migration in the L3 optic lobes

In addition to LPCs, the developing lamina is also populated by glia. Epithelial and marginal glia are positioned above and below photoreceptor growth cones (Figure 3A). These glia originate from glial precursor cell domains at the dorsal and ventral tips of the lamina and migrate tangentially into the developing lamina (Figure 3A) (Dearborn, 2004; Yoshida *et al*., 2005; Chen *et al*., 2016). When viewed in cross-section (Movie 5), we noticed that epithelial glia, situated above the photoreceptor growth cones (Figure 3A), were very motile and moved across photoreceptor growth cones and sometimes below to the level of marginal glia (Movie 5). Though they originate from the same domains, epithelial and marginal glia are distinct cell types (Chotard and Salecker, 2007; Edwards and Meinertzhagen, 2010; Edwards *et al*., 2012). While they express different molecular markers at later developmental stages, at L3 they have been distinguished solely by their by their relative positions on either side of photoreceptor growth cones (in fixed samples) (Chotard and Salecker, 2007; Edwards *et al*., 2012). In our L3 live imaging, lamina glial positions were not as stable as expected from previous descriptions, (Movie 5). We also observed glial migration towards the anterior side of the lamina from posterior positions (Movie 5), most likely a consequence of glial incorporation into young lamina columns. Epithelial and marginal glia have neuronal siblings, which develop into two neuron subtypes called lamina wide-field neurons 1 and 2 (Lawfs) (Chen *et al.*, 2016; Suzuki *et al.*, 2016). Lawfs are also born in the glial precursor cell domains and migrate tangentially but below the level of glia to incorporate into the deepest layers of the medulla (Chen *et al*., 2016). Since *gcm-GAL4* labels Lawfs as well as LPCs and lamina glia, we used it to express membrane-tagged GFP *(UAS-CD8::GFP)* to visualize glial and Lawf neuronal migration (Movie 6). Lawf migration was readily captured as described (Chen *et al*., 2016). However, the dense packing of labelled glia proved challenging for tracking these cells (Figure 3A and Movie 6). We therefore switched to a sparse labelling technique, called FlexAmp (Bertet *et al*., 2014) to induce stochastic and permanent expression of *myristoylated-GFP (myr-GFP)* in the glial precursor cell domains and thus progeny originating therein. Using *dpp*-gal4 to induce sparse labelling, GFP-positive cells were observed at the dorsal and ventral tips of the lamina and in the lobula plug, where *dpp-gal4* is expressed (Figure 3 and Movie 7). Furthermore, we could clearly track the migration of several glia originating from these domains into the lamina. These glia displayed many dynamic membrane protrusions (Movie 7), as inferred by others from fixed tissue (Poeck *et al*., 2001; Yoshida *et al*., 2005). Overall, we show that our live imaging protocol can be used to capture dynamic processes involved in cell migration during lamina development. We revealed cell behaviors that were not apparent from fixed tissue, including membrane protrusion dynamics during glial migration and epithelial and marginal glial incorporation into lamina columns, suggesting that these two glial cell types are not strictly separate until later in development.

### Eye disc eversion

During wL3 stage, the eye-antennal discs (EADs) undergo complex remodeling to give rise to several adult head structures and the head epidermis (reviewed by Kumar, 2018). One of the most prominent metamorphic events that primes the EADs to generate their corresponding adult appendages is disc evagination, which is subdivided into two discrete processes—elongation and eversion (reviewed by Gibson and Schubiger, 2001). The EADs are comprised of two epithelial layers, the columnar disc proper and a squamous epithelium called peripodial membrane. The peripodial membrane sits atop and is continuous with the disc proper (Figure 4), with interaction between the two described as vital for disc eversion (Milner *et al*., 1983). To date, studies focused on this dynamic event have been limited to fixed tissues (Gibson and Schubiger, 2001), largely attributed to absence of appropriate culturing and immobilization systems given anisotropic forces exerted by biological glues that likely affect morphogenesis (Kumar, 2018).

**Figure 4.**
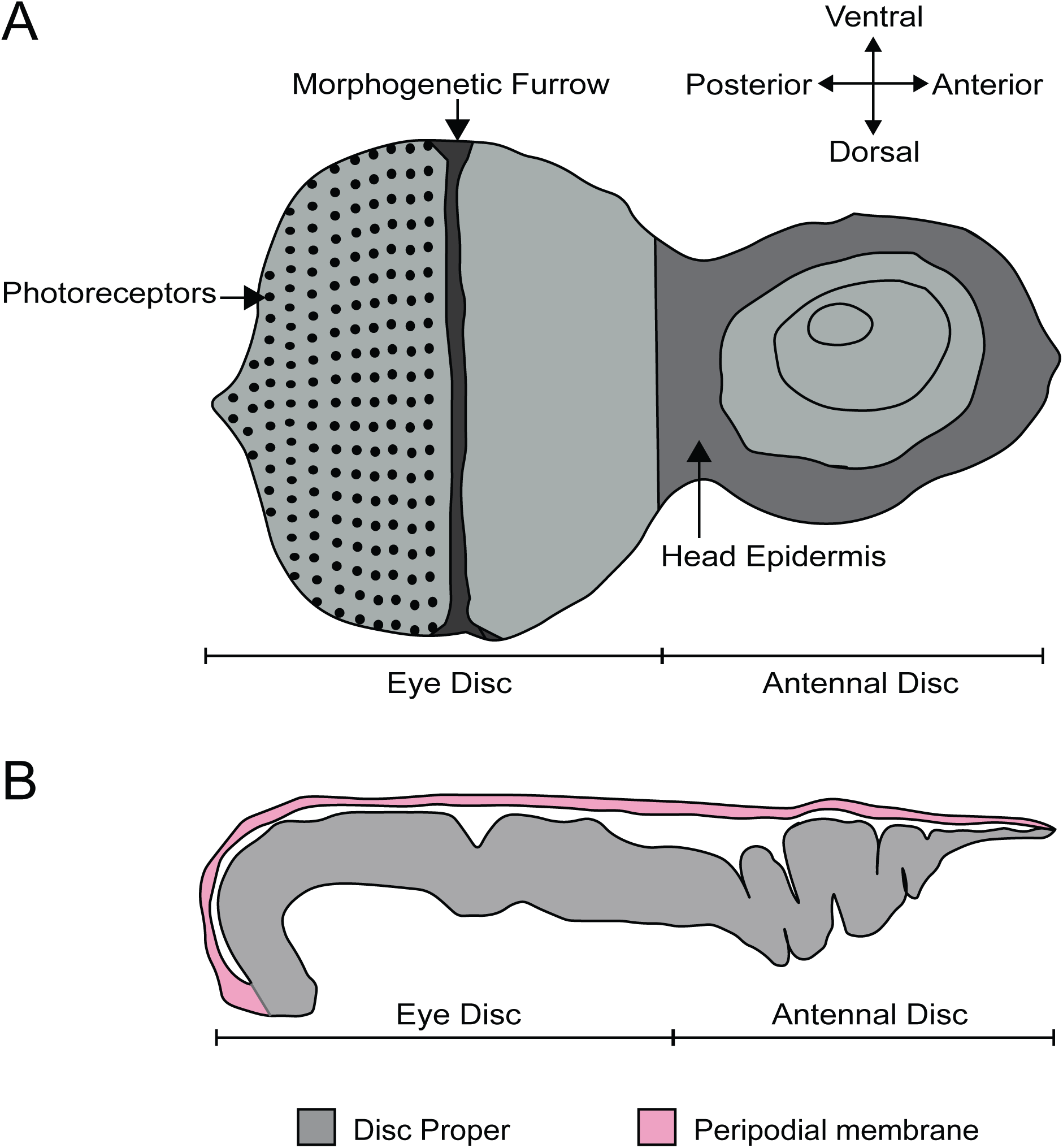
Eye-antennal disc structure. **A:** Schematic of L3 eye-antennal disc, adapted from Kumar, 2008. **B:** Cross-section of the eye-antennal disc. The columnar epithelium of the disc proper at the anterior end (antennal side) is folded whereas that of the eye disc is stretched and convex in shape. The thin peripodial membrane sits above and is continuous with the disc proper which constitutes the eye disc and the antennal disc.

We tested whether our culture system and immobilization technique could be used to capture EAD eversion. We dissected late L3 CNSs ubiquitously expressing a membrane-bound GFP (*ubi-GFP-CAAX*) with attached EADs. These explanted EADs immobilized in agarose underwent disc eversion within a period of five hours (Movie 8). The peripodial membrane appeared to contract and pull the eye disc proper towards the larval epidermis. The eye disc curled anteriorly taking on an ovular shape, at the same time the antennal disc was molded into a circular shape. This morphological change of the antennal discs is important to drive their movement outside the larval epidermis and then fusion to form the adult head epidermis (Milner *et al*., 1984). Since our culture system and immobilization technique recapitulated disc eversion events as observed in histological studies of cultured EADs carried out by others (Milner *et al*., 1983), it can be used to study and visualize in real time how the peripodial membrane affects disc eversion. For example, EADs in which the peripodial membrane is genetically or physically ablated can be imaged live to further analyze the mechanics of disc eversion.

### Stem cell maintenance in the adult testis

*Ex vivo* imaging can bypass many of the technical challenges associated with imaging adult tissues such as opaque cuticle and animal movement. To test whether our *ex vivo* imaging setup could be applied to adult tissue we focused on the *Drosophila* testis, a well-characterized model to study homeostatic mechanisms regulating stem cell behaviors in their intact microenvironment. The testis stem cell niche is composed of a cluster of quiescent stromal cells collectively known as the hub (Hardy *et al*., 1979). The hub is anchored at the apex of a blunt-ended coiled tube that forms the testis (Figure 5). Hub cells support two stem cell populations, germline stem cells (GSCs) and somatic cyst stem cells (CySCs). GSCs and CySCs are physically attached to the hub, and their self-renewal is maintained by hub-derived signals (Figure 5) (reviewed by Greenspan *et al.*, 2015). The two stem cell populations are easily distinguishable by morphology and position. GSCs are large and round cells that tightly associate with the hub whereas CySC have smaller nuclei located behind GSCs and extend a thin membrane projection between GSCs to contact the hub (Hardy *et al.*, 1979) (Figure 5).

**Figure 5.**
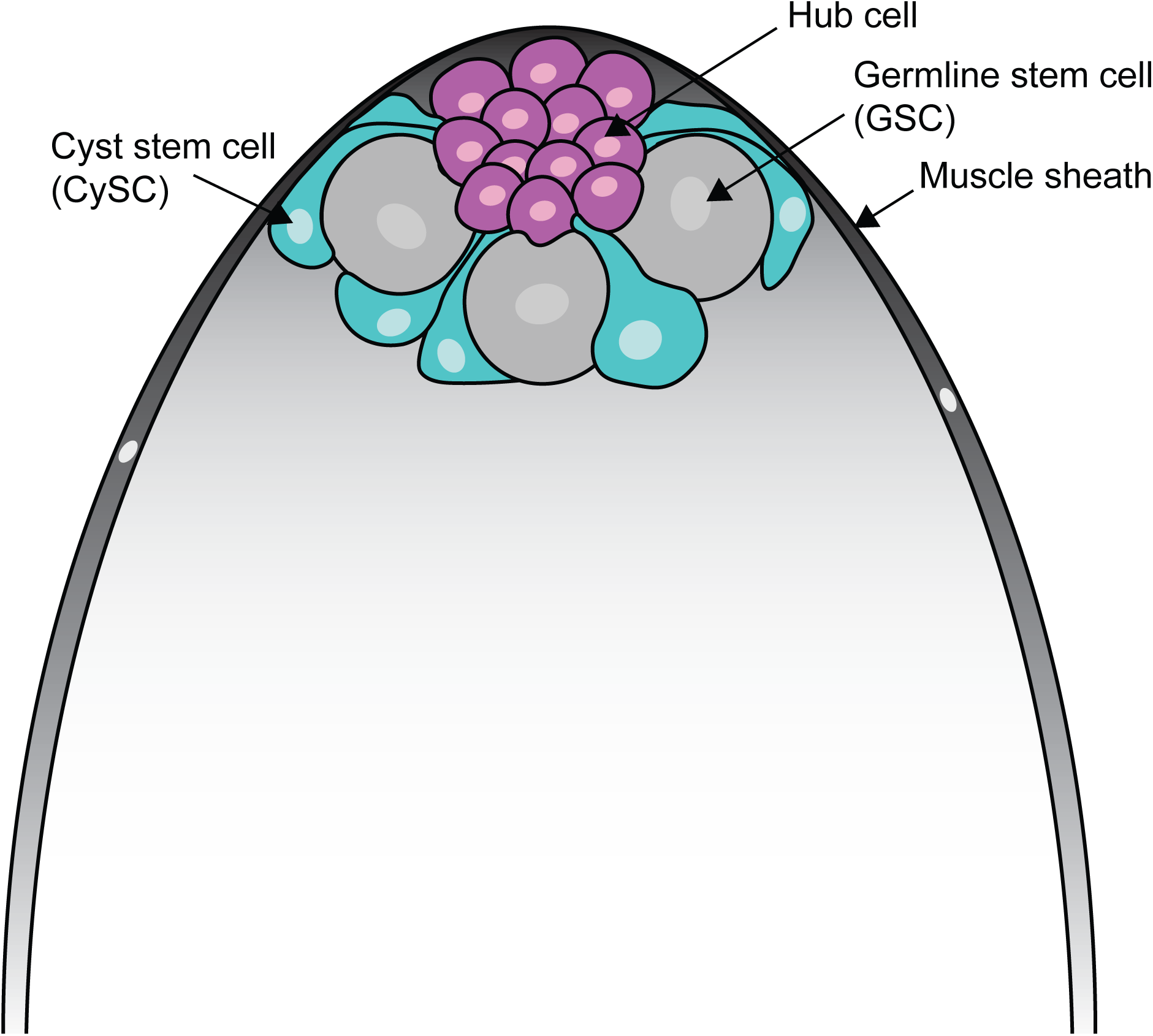
Schematic depiction of the spatial organization of stem and niche cells in the adult testis. The adult testis tissue is encased in a muscle sheath (grey). Anchored at the apical tip of the tissue is a cluster of small post-mitotic cells known as hub cells (pink), which constitute the stem cell niche. Two stem cell populations are in direct contact with the hub – germline stem cells (GSCs, grey) and cyst stem cells (CySCs, cyan). GSCs are large, round cells that closely associate with the hub. CySCs are located behind GSCs and extend thin membrane protrusions between GSCs to contact the hub.

Encased in a muscle sheath, the adult testis is a very contractile tissue that renders long-term live imaging technically challenging. Previous efforts to overcome this contractility typically involved weighing down the tissue onto the culture dish using cellulose membranes, teflon sheets or coverslips (Cheng and Hunt, 2009; Sheng and Matunis, 2011; Inaba *et al.*, 2015) or adhering the tissue to poly-L-lysine-coated coverslips (Lenhart and DiNardo, 2015; Greenspan and Matunis, 2017). These methods exert physical stress on the tissue during imaging. Here, we used our immobilization strategy to eliminate any non-specific effects of anisotropic forces exerted on the tissue. We used the somatic lineage-specific driver, *traffic jam (tj)-GAL4* to drive membrane targeted GFP (*UAS-CD8::GFP*), thus labelling the entire lineage including the CySCs and their progeny. This method did not eliminate muscle sheath contraction but as documented by others previously, not all testes are contractile, and these were chosen for imaging (Sheng and Matunis, 2011; Inaba *et al.*, 2015). Post-acquisition computational drift correction (Correct 3D Drift Plugin, ImageJ – see Methods) was sufficient to generate a stable movie for analysis for testes that displayed minor contractility. We observed multiple CySC divisions (Movie 9), during which CySC nuclei moved closer to the hub and rounded up. Unexpectedly, nuclear membrane labelling by CD8::GFP was apparent and persisted throughout CySC divisions, and we observed that this nuclear labelling could be reliably used to identify dividing CySCs (Movies 9 and 10). This observation suggests that CySCs divide with a closed or semi-closed nuclear division as has been reported in other *Drosophila* cells including embryos and germ cell meiosis (Church and Lin, 1982; Stafstrom and Staehelin, 1984; Debec and Marcaillou, 1997; Cheng *et al.*, 2011; Boettcher and Barral, 2013). This work demonstrates that our protocol can be adjusted to track cellular behaviors in contractile adult tissues.

### Conclusions

Live imaging enables novel insights into dynamic biological processes of different scales, from the subcellular to the multicellular. Here we detail a simple and inexpensive protocol for immobilizing explanted tissues in any precise orientation desired, which supports long-term live imaging with minimal physical stress. We validated our approach by visualizing dynamic processes previously described for L3 brains and adult testes and applied it to make novel observations: (1) multiple lamina columns undergo assembly together, (2) dynamic membrane protrusions and extensions of epithelial and marginal glia during migration, (3) fibers of quiescent neuroblasts are inherited by the GMC upon reactivation, and (4) the nuclear membrane of adult CySCs does not breakdown completely during mitosis.

Our protocol is amenable to customization with minimal effort and could be used for experimental approaches requiring temperature shifts (*e.g.* 29 **°**C for temperature-sensitive mutants or increased GAL4 activity), for approaches requiring acute drug treatment by combination with a flow perfusion apparatus (Williamson and Hiesinger, 2010), and to other species.

We note that this protocol is optimized for upright microscopes using a water-dipping objective. The upright set-up is useful for certain tissue orientations, but the protocol could be used with an inverted microscope using a glass-bottomed petri dish. For the latter, users should consider the objective working distance and the volume of agarose for immobilization, however. While we have not tested our protocol with an inverted set-up, the volume of agarose used to immobilize the tissue may need to be reduced or sample pushed closer to bottom to ensure sufficient proximity to the glass bottom for imaging.

Overall, this protocol offers a simple, inexpensive and versatile method to immobilize explanted tissues for long-term live imaging.

## Author Contributions

MPB and ARP designed and performed L3 experiments with *gcm-GAL4* and *dpp-GAL4* (MPB), and *ubi-GFP-CAAX* (ARP). MPB analysed the data, prepared the figures and movies and wrote the majority of the manuscript together with ARP. RC performed the L1 experiments, analysed the data, and prepared figures and movies; RC and RSN and wrote the corresponding text. ACY performed experiments related to the adult testis, analysed the data, prepared the figures and movies and wrote the corresponding text. RSN and MA contributed to writing and editing the manuscript. VMF designed and performed experiments with *His2av::EGFP* and dual-fluorophore labelling of the lamina, supervised the project, and contributed to writing and editing the manuscript.

## Acknowledgments

We are grateful to members of the Fernandes, Amoyel and Sousa-Nunes lab for critical comments on the manuscript.

## Figure Legends

**Movie** 1 **– Neuroblast divisions in the third instar brain lobe.** Time-lapse 2-photon imaging of wandering L3 (wL3) larval brain lobes. Example neuroblast and GMC divisions are highlighted; neuroblasts were identified by their large cell size. His2Av::eGFP marks chromosomes. Timescale displayed as hh:mm:ss, scale bar = 10 µm.

**Movie 2 – Neuroblasts reactivating from quiescence.** Time lapse 2-photon imaging of *grh*>*syn21-GFP-p10* labeled neuroblasts in late L1 VNC. Explants oriented laterally to best visualize the neuroblast fibers, present during quiescence. The movie starts 24 h after larval hatching, a time when most VNC neuroblasts are quiescent. Reactivation divisions were first observed at movie time 1 h 40 minutes. Compared with membrane-localized GFP, cytoplasmic GFP requires higher levels of expression to achieve similar brightness in fine cellular processes due to unfavorable surface area-to-volume ratios (Pfeiffer *et al*., 2012). Timescale displayed as hh:mm:ss, scale bar = 20 µm.

**Movie 3 – Basal fiber inheritance by firstborn post-reactivation ganglion mother cell.** Time lapse imaging of *grh*>*CD4::tdTomato* neuroblast in late L1 VNC undergoing first post-reactivation division. Movie captures fiber inheritance by the firstborn GMC. Timescale displayed as hh:mm:ss, scale bar = 10 µm.

**Movie 4 – Growth of the larval lamina.** Time-lapse 2-photon imaging of *E-Cad::GFP* (cyan), *27G05*>*mCherry* (magenta) demonstrating lamina growth. Brains were oriented to visualize the lamina in a lateral view. The lamina furrow, photoreceptors, and lamina columns were easily distinguishable using *E-Cad::GFP*; lamina precursors and neurons were labelled with *27G05*>*mCherry*. We observed that whilst the lamina increased in width during the course of the movie (∼18 hours) the position of the lamina furrow (asterisk) did not move significantly. Timescale displayed as hh:mm:ss, scale bar = 20 µm.

**Movie 5 – Incorporation of lamina precursor cells into lamina columns.** Time-lapse 2-photon imaging of gcm>nlsGFP. We saw multiple LPCs from the lamina assembly domain incorporate into columns. The positions of the lamina furrow, lamina pre-assembly domain, photoreceptor axon entry point, lamina columns and epithelial and marginal glia are marked. In the first half of the movie LPCs were tracked (colored dots) and seen to incorporate into multiple columns simultaneously. In the second half, epithelial and marginal were tracked (colored dots) with several re-locating to different positions. Note: Dots remain in the last observed position of a cell if we could not track it due to movement out of plane. We also observed the incorporation of an eg into the youngest lamina column. Timescale displayed as hh:mm:ss, scale bar = 20 µm.

**Movie 6 – Migration of lamina wide-field neurons and epithelial and marginal glia.** Time-lapse 2-photon imaging of *gcm*>*CD8::GFP* (expressed in Lawfs, epithelial and marginal glia and lamina precursors) captured the migration of Lawfs, and epithelial and marginal glia. The explant was oriented such that a cross-section of the lamina could be seen together with the dorsal arm of the lamina crescent. A maximum intensity projection is presented to account for movement in the z dimension. The video starts with Lawf migration marked by colored dots and the second half of the video shows epithelial and marginal glial migration, also marked by colored dots. Note: Dots remain in the last observed position of a cell if we could not track it due to movement out of plane. Timescale displayed as hh:mm:ss, scale bar = 20 µm.

**Movie 7 – Migration of epithelial and marginal glia.** Using *dpp*>*flexamp* to induce stochastic expression of myristoylated-GFP, time-lapse 2-photon imaging was employed to capture eg/mg migration. The brains were oriented such that a lateral view of the lamina could be seen. A maximum intensity projection is presented to account for movement in the z dimension. The lobula plug and glial progenitor domains of the OPC can be seen. Epithelial and marginal glia migrate towards the central region of the lamina (marked with arrows). Epithelial and marginal glia migrating are labelled on the right-hand side of the lamina and their migration path is indicated. A cropped zoom clearly shows filopodial-like projections in migrating epithelial and marginal glia. Timescale displayed as hh:mm:ss, scale bar = 20 µm.

**Movie 8 – Eye-antennal disc eversion in third instar larva.** Wandering L3 eye discs (attached to the underlying CNS) ubiquitously expressing membrane bound GFP (*ubi-GFP-CAAX*) to visualize eye-antennal disc eversion. As eversion begins, the peripodial epithelium is seen to contract (indicated by arrows) and pull the eye disc towards the antennal disc which becomes circular in shape. Disc eversion is completed within 5 hours. Timescale displayed as hh:mm:ss, scale bar = 40 µm.

**Movie 9 – Cyst stem cell (CySC) division in the adult testis stem cell niche.** A time-lapse movie of *tj*>*CD8::GFP*, which labels the somatic lineage. The hub is indicated by a magenta dot. CySCs are the first row of labelled cells around the hub. The CySC labelled with a blue dot undergoes a division to produce two daughter cells (blue and green dots). The nuclear membrane remains visible throughout mitosis of the CySC. Timescale displayed as hh:mm:ss, scale bar = 10 µm.

**Movie 10 – Nuclear envelope retention during CySC division.** A time-lapse movie of *tj*>*CD8::GFP*, which labels the somatic lineage. The hub is marked by a blue dot, a dividing CySC is marked by a green dot. During mitosis, the labelled CySC rounded up. Once again, the nuclear membrane could be clearly observed during mitosis and adopted a distinctive diamond shape. Note that the division occurs out of plane such that only one daughter is visible after the division. Timescale displayed as hh:mm:ss, scale bar = 20 µm.

## References

Aimon, S., Katsuki, T., Jia, T., Grosenick, L., Broxton, M., Deisseroth, K., et al. (2019). Fast near-whole-brain imaging in adult drosophila during responses to stimuli and behavior. PLoS Biol. 17, e2006732. doi: 10.1371/journal.pbio.2006732.

Akin, O., and Zipursky, S. L. (2016). Frazzled promotes growth cone attachment at the source of a Netrin gradient in the Drosophila visual system. Elife 5. doi: 10.7554/eLife.20762.

Aldaz, S., Escudero, L. M., and Freeman, M. (2010). Live imaging of Drosophila imaginal disc development. PNAS 107, 14217–14222. doi: 10.1073/pnas.1008623107.

Alexandre, P., Reugels, A. M., Barker, D., Blanc, E., and Clarke, J. D. W. (2010). Neurons derive from the more apical daughter in asymmetric divisions in the zebrafish neural tube. Nat. Neurosci. 13, 673–679. doi: 10.1038/nn.2547.

Baker, J., Theurkauf, W. E., and Schubiger, G. (1993). Dynamic changes in microtubule configuration correlate with nuclear migration in the preblastoderm Drosophila embryo. J. Cell Biol. 122, 113–121. doi: 10.1083/jcb.122.1.113.

Balaji, R., Bielmeier, C., Harz, H., Bates, J., Stadler, C., Hildebrand, A., et al. (2017). Calcium spikes, waves and oscillations in a large, patterned epithelial tissue. Sci. Rep. 7, 1–14. doi: 10.1038/srep42786.

Barlan, K., Cetera, M., and Horne-Badovinac, S. (2017). Fat2 and Lar Define a Basally Localized Planar Signaling System Controlling Collective Cell Migration. Dev. Cell 40, 467–477.e5. doi: 10.1016/j.devcel.2017.02.003.

Bell, D. M. (2017). Imaging morphogenesis. Phil. Trans. R. Soc. B. 372, 20150511. doi: 10.1098/rstb.2015.0511.

Bertet, C., Li, X., Erclik, T., Cavey, M., Wells, B., and Desplan, C. (2014). Temporal patterning of neuroblasts controls notch-mediated cell survival through regulation of hid or reaper. Cell 158, 1173–1186. doi: 10.1016/j.cell.2014.07.045.

Besson, C., Bernard, F., Corson, F., Rouault, H., Reynaud, E., Keder, A., et al. (2015). Planar cell polarity breaks the symmetry of PAR protein distribution prior to mitosis in Drosophila sensory organ precursor cells. Curr. Biol. 25, 1104–1110. doi: 10.1016/j.cub.2015.02.073.

Boettcher, B., and Barral, Y. (2013). The cell biology of open and closed mitosis. Nucleus 4, 160–165. doi: 10.4161/nucl.24676.

Bosveld, F., Bonnet, I., Guirao, B., Tlili, S., Wang, Z., Ambre, P., et al. (2012). Mechanical Control of Morphogenesis by Fat/Dachsous/Four-Jointed Planar Cell Polarity Pathway. Science (80-.). 336, 724–727. doi: 10.1126/science.1221071.

Britton, J. S., and Edgar, B. A. (1998). Environmental control of the cell cycle in Drosophila: nutrition activates mitotic and endoreplicative cells by distinct mechanisms. Development 125, 2149 LP–2158.

Cabernard, C., and Doe, C. Q. (2013). Live Imaging of Neuroblast Lineages within Intact Larval Brains in Drosophila. Cold Spring Harb. Protoc., 970–977. doi: 10.1101/pdb.prot078162.

Cavey, M., and Lecuit, T. (2008). Imaging Cellular and Molecular Dynamics in Live Embryos Using Fluorescent Proteins. Methods Mol. Biol. 420, 219–238. Available at: 10.1007/978-1-59745-583-1_13.

Chell, J. M., and Brand, A. H. (2010). Nutrition-responsive glia control exit of neural stem cells from quiescence. Cell 143, 1161–1173. doi: 10.1016/j.cell.2010.12.007.

Chen, Z., Del Valle Rodriguez, A., Li, X., Erclik, T., Fernandes, V. M., and Desplan, C. (2016). A Unique Class of Neural Progenitors in the Drosophila Optic Lobe Generates Both Migrating Neurons and Glia. Cell Rep. 15, 774–786. doi: 10.1016/J.CELREP.2016.03.061.

Cheng, J., and Hunt, A. J. (2009). Time-lapse live imaging of stem cells in drosophila testis. Curr. Protoc. Stem Cell Biol. 0331, 1–9. doi: 10.1002/9780470151808.sc02e02s11.

Cheng, J., Tiyaboonchai, A., Yamashita, Y. M., and Hunt, A. J. (2011). Asymmetric division of cyst stem cells in Drosophila testis is ensured by anaphase spindle repositioning. Development 138, 831–837. doi: 10.1242/dev.057901.

Chotard, C., and Salecker, I. (2007). Glial cell development and function in the Drosophila visual system. Neuron Glia Biol. 3, 17–25. doi: 10.1017/S1740925x07000592.

Church, K., and Lin, H. P. (1982). Meiosis in Drosophila melanogaster. II. The prometaphase-I kinetochore microtubule bundle and kinetochore orientation in males. J. Cell Biol. 93, 365–373. doi: 10.1083/jcb.93.2.365.

Dearborn, R. (2004). An axon scaffold induced by retinal axons directs glia to destinations in the Drosophila optic lobe. Development 131, 2291–2303. doi: 10.1242/dev.01111.

Debec, A., and Marcaillou, C. (1997). Structural alterations of the mitotic apparatus induced by the heat shock response in Drosophila cells. Biol. cell 89, 67—78. doi: 10.1016/s0248-4900(99)80082-3.

Distel, M., and Koster, R. W. (2007). In Vivo Time-Lapse Imaging of Zebrafish Embryonic Development. Cold Spring Harb. Protoc. 2007, pdb.prot4816-pdb.prot4816. doi: 10.1101/pdb.prot4816.

Dye, N. A., Popović, M., Spannl, S., Etournay, R., Kainmüller, D., Ghosh, S., et al. (2017). Cell dynamics underlying oriented growth of the drosophila wing imaginal disc. Development 144, 4406–4421. doi: 10.1242/dev.155069.

Echalier, G. (1997). “Composition of the Body Fluid of Drosophila and the Design of Culture Media for Drosophila Cells” in Drosophila Cells in Culture. ed. G. Echalier (Academic Press) 1–67. doi: 10.1016/B978-012229460-0/50002-6.

Edwards, T. N., and Meinertzhagen, I. A. (2010). The functional organisation of glia in the adult brain of Drosophila and other insects. Prog. Neurobiol. 90, 471–497. doi: 10.1016/j.pneurobio.2010.01.001.

Edwards, T. N., Nuschke, A. C., Nern, A., and Meinertzhagen, I. A. (2012). Organization and Metamorphosis of Glia in the Drosophila Visual System. 2085, 2067–2085. doi: 10.1002/cne.23071.

Farhadifar, R., Ro, J., Aigouy, B., and Eaton, S. (2007). The Influence of Cell Mechanics, Cell-Cell Interactions, and Proliferation on Epithelial Packing. Curr. Biol. 17, 2095–2104. doi: 10.1016/j.cub.2007.11.049.

Fernandes, V. M., Chen, Z., Rossi, A. M., Zipfel, J., and Desplan, C. (2017). Glia relay differentiation cues to coordinate neuronal development in *Drosophila*. Science (80-.). 357, 886–891. doi: 10.1126/science.aan3174.

Fiala, A., Spall, T., Eisermann, B., Sachse, S., Devaud, J., Buchner, E., et al. (2002). Genetically Expressed Cameleon in Drosophila melanogaster Is Used to Visualize Olfactory Information in Projection Neurons. Curr. Biol. 12, 1877–1884. doi: 10.1016/s0960-9822(02)01239-3.

Foe, V. E., and Alberts, B. M. (1983). Studies of nuclear and cytoplasmic behaviour during the five mitotic cycles that precede gastrulation in Drosophila embryogenesis. J. Cell Sci. 61, 31–70.

Ghannad-Rezaie, M., Wang, X., Mishra, B., Collins, C., and Chronis, N. (2012). Microfluidic chips for in vivo imaging of cellular responses to neural injury in Drosophila larvae. PLoS One 7, e29869. doi: 10.1371/journal.pone.0029869.

Gibson, M. C., and Schubiger, G. (2001). Drosophila peripodial cells, more than meets the eye? BioEssays 23, 691–697.

Greenspan, L. J., de Cuevas, M., and Matunis, E. (2015). Genetics of Gonadal Stem Cell Renewal. Annu. Rev. Cell Dev. Biol. 31, 291–315. doi: 10.1146/annurev-cellbio-100913-013344.

Greenspan, L. J., and Matunis, E. L. (2017). “Live imaging of the drosophila testis stem cell niche,” in Methods in Molecular Biology (Humana Press Inc.), 63–74. doi: 10.1007/978-1-4939-4017-2_4.

Hansen, D. V., Lui, J. H., Parker, P. R. L., and Kriegstein, A. R. (2010). Neurogenic radial glia in the outer subventricular zone of human neocortex. Nature 464, 554–561. doi: 10.1038/nature08845.

Hardy, R. W., Tokuyasu, K. T., Lindsley, D. L., and Garavito, M. (1979). The germinal proliferation center in the testis of Drosophila melanogaster. J. Ultrasructure Res. 69, 180–190. doi: 10.1016/S0022-5320(79)90108-4.

Heemskerk, I., Lecuit, T., and LeGoff, L. (2014). Dynamic clonal analysis based on chronic in vivo imaging allows multiscale quantification of growth in the Drosophila wing disc. Development 141, 2339–2348. doi: 10.1242/dev.109264.

Hellerman, M. B., Choe, R. H., and Johnson, R. I. (2015). Live-imaging of the Drosophila pupal eye. J. Vis. Exp., 1–9. doi: 10.3791/52120.

Homem, C. C. F., Reichardt, I., Berger, C., Lendl, T., and Knoblich, J. A. (2013). Long-term live cell imaging and automated 4D analysis of Drosophila neuroblast lineages. PLoS One 8, 1–10. doi: 10.1371/journal.pone.0079588.

Huang, J., Zhou, W., Dong, W., Watson, A. M., and Hong, Y. (2009). Directed, efficient, and versatile modifications of the Drosophila genome by genomic engineering. Proc. Natl. Acad. Sci. U. S. A. 106, 8284–8289. doi: 10.1073/pnas.0900641106.

Huang, Z., and Kunes, S. (1996). Hedgehog, Transmitted along Retinal Axons, Triggers Neurogenesis in the Developing Visual Centers of the Drosophila Brain. Cell 86, 411–422. doi: 10.1016/S0092-8674(00)80114-2

Huang, Z., and Kunes, S. (1998a). Signals transmitted along retinal axons in Drosophila : Hedgehog signal reception and the cell circuitry of lamina cartridge assembly. Development 125, 3753–3764.

Huang, Z., Shilo, B., and Kunes, S. (1998b). A Retinal Axon Fascicle Uses Spitz, an EGF Receptor Ligand, to Construct a Synaptic Cartridge in the Brain of Drosophila. Cell 95, 693–703. doi: 10.1016/S0092-8674(00)81639-6.

Huisken, J., and Stainier, D. Y. R. (2009). Selective plane illumination microscopy techniques in developmental biology. Development 136, 1963–1975. doi: 10.1242/dev.022426.

Ichikawa, T., Nakazato, K., Keller, P. J., Kajiura-Kobayashi, H., Stelzer, E. H. K., Mochizuki, A., et al. (2014). Live imaging and quantitative analysis of gastrulation in mouse embryos using light-sheet microscopy and 3D tracking tools. Nat. Protoc. 9, 575–585. doi: 10.1038/nprot.2014.035.

Inaba, M., Buszczak, M., and Yamashita, Y. M. (2015). Nanotubes mediate niche-stem-cell signalling in the Drosophila testis. Nature 523, 329–332. doi: 10.1038/nature14602.

Jacinto, A., Wood, W., Balayo, T., Turmaine, M., Martinez-arias, A., and Martin, P. (2000). Dynamic actin-based epithelial adhesion and cell matching during Drosophila dorsal closure. Curr. Biol. 10, 1420–1426. doi: 10.1016/s0960-9822(00)00796-x.

Kaltschmidt, J. A., Davidson, C. M., Brown, N. H., and Brand, A. H. (2000). Rotation and asymmetry of the mitotic spindle direct asymmetric cell division in the developing central nervous system. Nat. Cell Biol. 2, 7–12. doi: 10.1038/71323.

Kiehart, D. P., Galbraith, C. G., Edwards, K. A., Rickoll, W. L., and Montague, R. A. (2000). Multiple Forces Contribute to Cell Sheet Morphogenesis for Dorsal Closure. J. Cell Biol. 149, 471–490. doi: 10.1083/jcb.149.2.47.

Konno, D., Shioi, G., Shitamukai, A., Mori, A., Kiyonari, H., Miyata, T., et al. (2008). Neuroepithelial progenitors undergo LGN-dependent planar divisions to maintain self-renewability during mammalian neurogenesis. Nat. Cell Biol. 10, 93–101. doi: 10.1038/ncb1673.

Kumar, J. P. (2018). The Fly Eye : Through the Looking Glass. Dev. Dyn. 247, 111–123. doi: 10.1002/dvdy.24585.

Lee, C., Andersen, R. O., Cabernard, C., Manning, L., Tran, K. D., Lanskey, M. J., et al. (2006). Drosophila Aurora-A kinase inhibits neuroblast self-renewal by regulating aPKC / Numb cortical polarity and spindle orientation. Genes Dev 20, 3464–3474. doi: 10.1101/gad.1489406.Neuroblast.

Lenhart, K. F., and DiNardo, S. (2015). Somatic cell encystment promotes abscission in germline stem cells following a regulated block in cytokinesis. Dev. Cell 34, 192–205. doi: 10.1016/j.devcel.2015.05.003.

Lerit, D. A., Plevock, K. M., and Rusan, N. M. (2014). Live imaging of Drosophila larval neuroblasts. J. Vis. Exp., 1–12. doi: 10.3791/51756.

Mao, Y., Tournier, A. L., Bates, P. A., Gale, J. E., Tapon, N., and Thompson, B. J. (2011). Planar polarization of the atypical myosin Dachs orients cell divisions in Drosophila. Genes Dev. 25, 131–136. doi: 10.1101/gad.610511.GENES.

Martin, J. L., Sanders, E. N., Moreno-roman, P., Ann, L., Koyama, J., Balachandra, S., et al. (2018). Long-term live imaging of the Drosophila adult midgut reveals real-time dynamics of division, differentiation and loss. Elife 7, e36248. doi: 10.7554/eLife.36248.

Milner, M. J., Bleasby, A. J., and Pyott, A. (1983). The Role of the Peripodial Membrane in the Morphogenesis of the Eye-Antennal Disc of Drosophila melanogaster. Roux’s Arch Dev Biol 192, 164–170.

Milner, M. J., Bleasby, A. J., and Pyott, A. (1984). Cell interactions during the fusion in vitro of Drosophila eye-antennal imaginal discs. Roux’s Arch Dev Biol 193, 406–413.

Miyata, T., Kawaguchi, A., Okano, H., and Ogawa, M. (2001). Asymmetric inheritance of radial glial fibers by cortical neurons. Neuron 31, 727–741. doi: 10.1016/S0896-6273(01)00420-2.

Nickerson, P. E., Ronellenfitch, K. M., Csuzdi, N. F., Boyd, J. D., Howard, P. L., Delaney, K. R., et al. (2013). Live imaging and analysis of postnatal mouse retinal development. BMC Dev. Biol. 13, 1–17. doi: 10.1186/1471-213X-13-24.

Özel, M. N., Langen, M., Hassan, B. A., and Hiesenger, R. P. (2015). Filopodial dynamics and growth cone stabilization in Drosophila visual circuit development. Elife 4. doi: 10.7554/eLife.10721.

Parslow, A., Cardona, A., and Bryson-Richardson, R. J. (2014). Sample drift correction following 4D confocal time-lapse Imaging. J. Vis. Exp. doi: 10.3791/51086.

Parton, R. M., Vallés, A. M., Dobbie, I. M., and Davi, I. (2010). Live cell imaging in Drosophila melanogaster. Cold Spring Harb. Protoc. 5, pdb.top75. doi: 10.1101/pdb.top75.

Pfeiffer, B. D., Truman, J. W., and Rubin, G. M. (2012). Using translational enhancers to increase transgene expression in Drosophila. Proc. Natl. Acad. Sci. U. S. A. 109, 6626–6631. doi: 10.1073/pnas.1204520109.

Poeck, B., Fischer, S., Gunning, D., Zipursky, S. L., and Salecker, I. (2001). Glial cells mediate target layer selection of retinal axons in the developing visual system of Drosophila. Neuron 29, 99–113. doi: 10.1016/S0896-6273(01)00183-0.

Prasad, M., Wang, X., He, L., Cai, D., and Montell, D. J. (2015). Border cell migration: A model system for live imaging and genetic analysis of collective cell movement. Methods Mol. Biol. 1328, 89–97. doi: 10.1007/978-1-4939-2851-4_6.

Rabinovich, D., Mayseless, O., and Schuldiner, O. (2015). Long term ex vivo culturing of Drosophila brain as a method to live image pupal brains : insights into the cellular mechanisms of neuronal remodeling. Front. Cell. Neurosci. 9. doi: 10.3389/fncel.2015.00327.

Restrepo, S., Zartman, J. J., and Basler, K. (2016). “Cultivation and Live Imaging of Drosophila Imaginal Discs,” in Drosophila: Methods and Protocols, Methods in Molecular Biology, ed. C. Dahmann (New York: Springer Science+Business Media), 203–213. doi: 10.1007/978-1-4939-6371-3_11.

Ritsma, L., Ellenbroek, S. I. J., Zomer, A., Snippert, H. J., De Sauvage, F. J., Simons, B. D., et al. (2014). Intestinal crypt homeostasis revealed at single-stem-cell level by in vivo live imaging. Nature 507, 362–365. doi: 10.1038/nature12972.

Rossi, A. M., and Fernandes, V. M. (2018). Wrapping Glial Morphogenesis and Signaling Control the Timing and Pattern of Neuronal Differentiation in the Drosophila Lamina. J. Exp. Neurosci. 12. doi: 10.1177/1179069518759294.

Sanchez-Corrales, Y. E., Blanchard, G. B., and Röper, K. (2018). Radially patterned cell behaviours during tube budding from an epithelium. Elife 7. doi: 10.7554/eLife.35717.

Sano, H., Renault, A. D., and Lehmann, R. (2005). Control of lateral migration and germ cell elimination by the Drosophila melanogaster lipid phosphate phosphatases Wunen and Wunen 2. J. Cell Biol. 171, 675–683. doi: 10.1083/jcb.200506038.

Sato, M., Suzuki, T., and Nakai, Y. (2013). Waves of differentiation in the fly visual system. Dev. Biol. 380, 1–11. doi: 10.1016/j.ydbio.2013.04.007.

Savoian, M. S., and Rieder, C. L. (2002). Mitosis in primary cultures of Drosophila melanogaster larval neuroblasts. J. Cell Sci. 115, 3061–3072.

Seelig, J. D., Chiappe, M. E., Lott, G. K., Dutta, A., Osborne, J. E., Reiser, M. B., et al. (2010). Two-photon calcium imaging from head-fixed Drosophila during optomotor walking behavior. Nat. Methods 7, 535–540. doi: 10.1038/nmeth.1468.

Selleck, S. B., and Steller, H. (1991). The Influence of Retinal Innervation on Neurogenesis in the First Optic Ganglion of Drosophila. Neuron 6, 83–99. doi: 10.1016/0896-6273(91)90124-i.

Sheng, X. R., and Matunis, E. (2011). Live imaging of the Drosophila spermatogonial stem cell niche reveals novel mechanisms regulating germline stem cell output. 3376, 3367–3376. doi: 10.1242/dev.065797.

Shitamukai, A., Konno, D., and Matsuzaki, F. (2011). Oblique radial glial divisions in the developing mouse neocortex induce self-renewing progenitors outside the germinal zone that resemble primate outer subventricular zone progenitors. J. Neurosci. 31, 3683–3695. doi: 10.1523/JNEUROSCI.4773-10.2011.

Siller, K. H., and Doe, C. Q. (2008). Lis1 / dynactin regulates metaphase spindle orientation in Drosophila neuroblasts. Dev. Biol. 319, 1–9. doi: 10.1016/j.ydbio.2008.03.018.

Siller, K. H., Serr, M., Steward, R., Hays, T. S., and Doe, C. Q. (2005). Live Imaging of Drosophila Brain Neuroblasts Reveals a Role for Lis1 / Dynactin in Spindle Assembly and Mitotic Checkpoint Control □. Mol. Biol. Cell 16, 5127–5140. doi: 10.1091/mbc.E05.

Sousa-Nunes, R., and Hirth, F. (2016). “Stem Cells and Asymmetric Cell Division,” in Regenerative Medicine - from Protocol to Patient: 1. Biology of Tissue Regeneration, ed. G. Steinhoff (Cham: Springer International Publishing), 87–121. doi: 10.1007/978-3-319-27583-3_3.

Sousa-Nunes, R., Yee, L. L., and Gould, A. P. (2011). Fat cells reactivate quiescent neuroblasts via TOR and glial insulin relays in Drosophila. Nature 471, 508–513. doi: 10.1038/nature09867.

Speder, P., and Brand, A. H. (2014). Article Gap Junction Proteins in the Blood-Brain Barrier Control Nutrient-Dependent Reactivation of Drosophila Neural Stem Cells. Dev. Cell 30, 309–321. doi: 10.1016/j.devcel.2014.05.021.

Stafstrom, J. P., and Staehelin, L. A. (1984). Dynamics of the nuclear envelope and of nuclear pore complexes during mitosis in the Drosophila embryo. Eur. J. Cell Biol. 34, 179–189.

Sugie, A., Umetsu, D., Yasugi, T., Fischbach, K. F., and Tabata, T. (2010). Recognition of pre- and postsynaptic neurons via nephrin/NEPH1 homologs is a basis for the formation of the Drosophila retinotopic map. Development 137, 3303–3313. doi: 10.1242/dev.047332.

Suzuki, T., Hasegawa, E., Nakai, Y., Kaido, M., Takayama, R., and Sato, M. (2016). Formation of Neuronal Circuits by Interactions between Neuronal Populations Derived from Different Origins in the Drosophila Visual Center. Cell Rep. 15, 499–509. doi: 10.1016/j.celrep.2016.03.056.

Tan, L., Zhang, K. X., Pecot, M. Y., Nagarkar-Jaiswal, S., Lee, P. T., Takemura, S. Y., et al. (2015). Ig Superfamily Ligand and Receptor Pairs Expressed in Synaptic Partners in Drosophila. Cell 163, 1756–1769. doi: 10.1016/j.cell.2015.11.021.

Truman, J. W., and Bate, M. (1988). Spatial and temporal patterns of neurogenesis in the central nervous system of Drosophila melanogaster. Dev. Biol. 125, 145–157. doi: 10.1016/0012-1606(88)90067-X.

Tsao, C. K., Ku, H. Y., Lee, Y. M., Huang, Y. F., and Sun, Y. H. (2016). Long term ex vivo culture and live imaging of Drosophila larval imaginal discs. PLoS One 11, e0163744. doi: 10.1371/journal.pone.0163744.

Umetsu, D., Murakami, S., Sato, M., and Tabata, T. (2006). The highly ordered assembly of retinal axons and their synaptic partners is regulated by Hedgehog / Single-minded in the Drosophila visual system. 791–800. doi: 10.1242/dev.02253.

Valcourt, J. R., Lemons, J. M. S., Haley, E. M., Kojima, M., Demuren, O. O., and Coller, H. A. (2012). Staying alive: Metabolic adaptations to quiescence. Cell Cycle 11, 1680–1696. doi: 10.4161/cc.19879.

Weissman, T., Noctor, S. C., Clinton, B. K., Honig, L. S., and Kriegstein, A. R. (2003). Neurogenic radial glial cells in reptile, rodent and human: from mitosis to migration. Cereb. Cortex 13, 550–559. doi: 10.1093/cercor/13.6.550.

Williamson, W. R., and Hiesinger, P. R. (2010). Preparation of Developing and Adult Drosophila Brains and Retinae for Live Imaging. J. Vis. Exp. 37, e1936. doi: 10.3791/1936.

Wyss, C. (1982). Ecdysterone, Insulin and Fly Extract needed for the proliferation of normal Drosophila cells in defined medium. Exp. Cell Res. 139, 297–307.

Yoshida, S., Soustelle, L., Giangrande, A., Umetsu, D., Murakami, S., Yasugi, T., et al. (2005). DPP signaling controls development of the lamina glia required forretinal axon targeting in the visual system of Drosophila. Development 132, 4587–4598. doi: 10.1242/dev.02040.

Zartman, J., Restrepo, S., Basler, K., Zartman, J., Restrepo, S., and Basler, K. (2013). A high-throughput template for optimizing Drosophila organ culture with response-surface methods. Development 2848, 667–674. doi: 10.1242/dev.098921.

