## Supplementary Material for "An immobilization technique for long-term time-lapse imaging of explanted *Drosophila* tissues"

### 1 Supplementary Data

**Printable protocol for culturing *Drosophila* central nervous system**
The following protocol is designed for culturing the *Drosophila* third larval instar central nervous system (CNS). We also demonstrated how this protocol can be applied to other tissues and developmental stages. Attached to this section is a schematic, outlining the steps involved.

**Agarose preparation**

1. Dissolve the appropriate amounts of low gelling temperature agarose (Sigma A9414) in sterile water to make a 2% solution. (Note: 2g in 98mL of sterile water lasts approximately 1 month stored at 4 °C). Melt the agarose using a microwave on high power, stirring every 30 seconds until the solution becomes clear.
2. Allow the agarose solution to solidify in glass bottles or 50mL falcon tubes. Once solid, cut into cubes (~ $2cm^3$ each) with a sterile instrument and place in distilled water at 4 °C. Replace the distilled water with fresh distilled water each day for three days – a necessary step to de-ionize the agarose.

**Culture medium**

3. Prepare culture medium in sterile conditions next to a flame. 50mL of medium is prepared from: 44.4mL Schneider's medium, 125µL human insulin (Sigma: 19278), 500µL Penicillin-Streptomycin (Sigma P4333), and 5mL FBS. 50-100mL lasts approximately 1 month stored at 4 °C.

**Dissection and Sample Immobilization**

4. To make the immobilization solution, heat 800µL of culture medium in 1.5mL Eppendorf tubes to 42 °C in a programmable heating block. Then add 200µL of 2% agarose (melted in the microwave) to each Eppendorf tube and mix well by pipetting or using a vortex.
5. Next, lower the temperature of the heating block to 34 °C and allow the immobilization solution in the Eppendorf tubes to cool to this temperature. Begin dissections during this time.
6. *Drosophila* explants must be dissected in culture medium. Take care to remove any unwanted tissues and leave the tissue of interested unscathed.
7. Once the 1mL agarose solution cools to 34 °C, pipette into an untreated 35x10mm suspension dish with lid and vent (Thermo 171099) so that it covers the whole dish in an even layer.
8. Transfer *Drosophila* explant to the periphery of the dish using forceps, taking care to transfer a minimal amount of culture medium to the agarose. Maneuver the sample to the center of the dish and orient accordingly. Use forceps to maneuver the tissue, moving the viscous agarose rather than the tissue itself. Complete all movements and orientations of the tissue within 5 mins of placing the brain in the agarose so as not to disrupt its setting. It is important that the tissue is at the center of the dish so that the immersion lens does not hit the wall of the petri dish.
9. After orienting, leave the agarose to set for at least 10 minutes.

10. Once the agarose has set, add 4mL of cold culture medium in a drop-wise manner, pipetting against the edge of the dish. This ensures that the medium does not go below the agarose layer.

**Troubleshooting**

*Bubbles in agarose/media.*
The presence of bubbles in the agarose can make it difficult to orient explants. It is important to remove bubbles with a pipette tip to ensure standard orientation.

*Early death of explant.*
When dissecting larval brains, it is imperative that the brain is not damaged. Any tear or loss of brain region causes an opening in the glial layer that forms the blood brain barrier. This can lead to ionic shock, resulting in death. Dying samples are apparent as fluorophores accumulate in bright puncta and cellular movements and divisions slow dramatically.

If the sample is dying sooner than expected, it could be that the media is contaminated. Check the culture medium to see if it is clear or cloudy. If cloudy, throw away immediately and make another aliquot.

*Moving samples.*
Explants can move for a variety of reasons. Firstly, in the case of the larval brain, the esophagus is attached to the mouth hooks. This contracts periodically even after dissection leading to movements of the entire sample. Immobilizing the sample can be achieved by crushing the esophagus with forceps or removing the it completely.

Sample drift can also be caused by high laser power. High laser energy can heat up and melt the agarose in which the explant sits, causing the sample to sink. Be mindful of how much dissection medium is transferred to the petri dish as this may locally dilute the agarose and allow sample drift.

*Difficulty orienting explant.*
When positioning the explant in a specific orientation it can be challenging to set it in the desired orientation. To combat this, when the 0.4% agarose is transferred to the culture dish, allow it to cool for 2-3 mins. This allows the agarose to become more viscous, and thus allows easier changes to the explant orientation.

### 2 Supplementary Figure 1: Step-by-Step protocol for sample immobilization in agarose for imaging

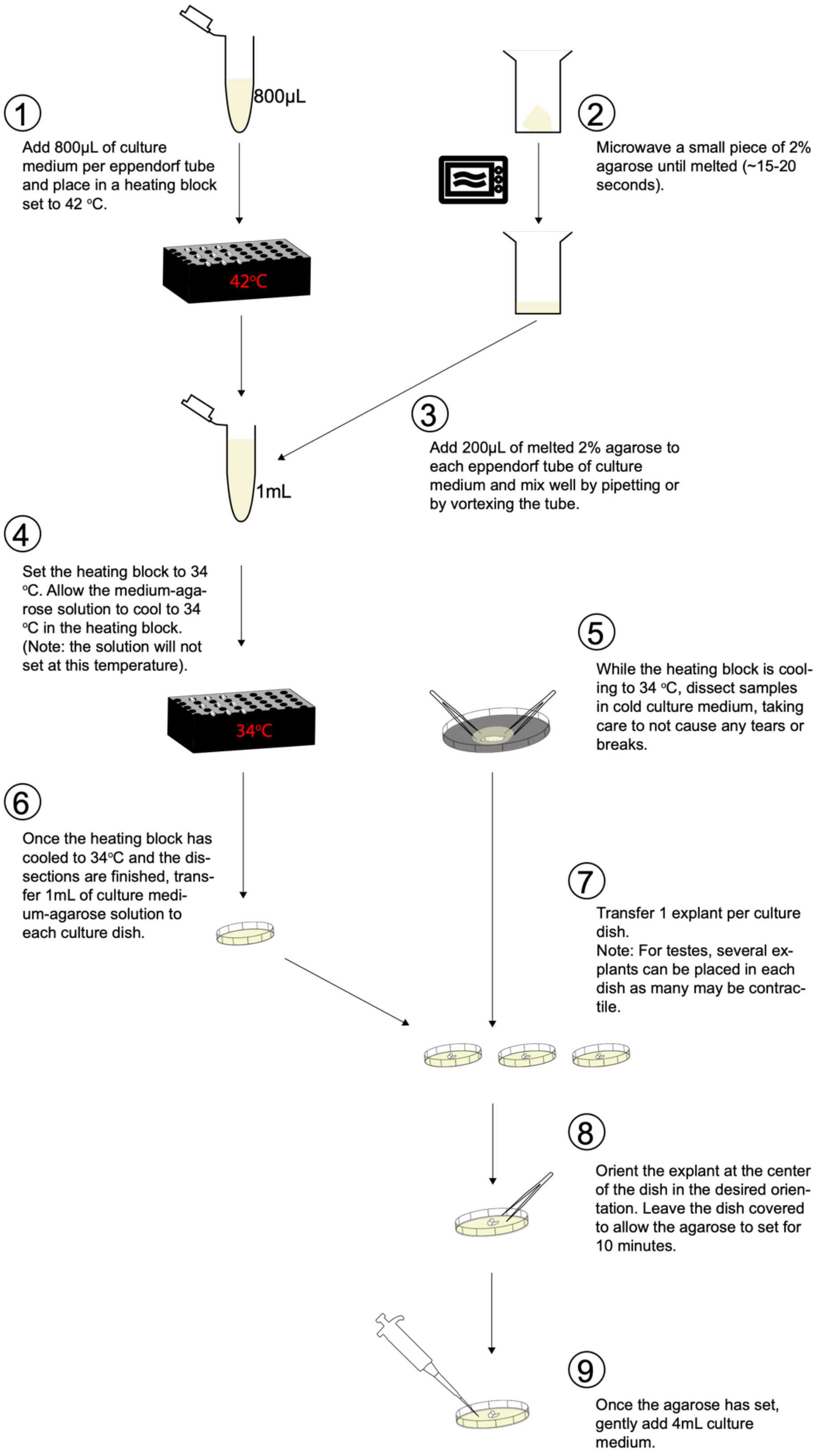
